# Inducible Nitric Oxide Synthase deficiency leads to early demyelination by altering the balance between pro- and anti-inflammatory responses against Murine-β-Coronavirus

**DOI:** 10.1101/2022.02.08.479662

**Authors:** Mithila Kamble, Fareeha Saadi, Saurav Kumar, Bhaskar Saha, Jayasri Das Sarma

## Abstract

The neurological disease Multiple sclerosis (MS) is characterized by neuroinflammation and demyelination orchestrated by the activated glial cells, the CNS infiltrating leukocytes, and their reciprocal interaction through inflammatory signals. Inducible nitric oxide synthase (iNOS), an enzyme that catalyzes sustained nitric oxide production in response to an inflammatory stimulus, is a pro-inflammatory marker expressed particularly by the microglia/macrophages (MG/Mφ) during neuroinflammation. In MS, iNOS has been reportedly associated with the disease pathology; however, studies dissecting its role in the underlying mechanisms, specifically demyelination, are limited. Therefore, we studied the role of iNOS in a recombinant beta-coronavirus-MHV-RSA59-induced neuroinflammation, which is a prototypic animal model used to investigate the pathological hallmarks of MS, neuroinflammatory demyelination, and axonal degeneration. During the acute phase of infection with RSA59, wildtype C57BL/6 (WT) mice had significantly upregulated iNOS expression in macrophages, natural killer cells, and natural killer T cells suggesting a role for iNOS in RSA59-induced neuroinflammation. Studies comparing RSA59-infected WT and iNOS-deficient mice revealed that iNOS deficiency aggravated the disease with increased CNS infiltration of macrophages and neutrophils and enhanced mortality. As early as 9-10 days after the infection, the CNS of iNOS-deficient mice had substantially higher demyelination marked with morphologically defined MG/Mφ in the demyelinating regions. Transcript analysis confirmed the significant upregulation of type2 macrophage (M2) markers-Arginase 1, CD206, and TREM2-in the CNS of iNOS-deficient mice. Corroborating to the phenotype, the iNOS-deficient mice showed a significantly higher expression of TGFβ-an anti-inflammatory cytokine- and increased T regulatory (Treg) cell infiltration, indicating an anti-inflammatory milieu established early after the infection. These observations highlight a protective role of iNOS in virus-induced neuroinflammation whereas its absence leads to MG/Mφ polarization towards a phenotype that may be involved in the exacerbated demyelination pathology.

**Author summary:** Contrary to the reported pathogenic role of inducible nitric oxide synthase (iNOS) in multiple sclerosis and related autoimmune animal models, we show that the mice deficient in iNOS show an exacerbated disease with accelerated demyelination accompanied by heightened production of an anti-inflammatory and phagocytic markers and more numbers of Tregs in a mouse model of a recombinant mouse hepatitis virus RSA59 infection. Therefore, iNOS may play protective and regulatory roles in this beta-coronavirus infection.

## Background

Multiple sclerosis (MS) is a neurological disease that is common among young adults and remains the most widely studied neuroinflammatory and demyelinating disorder of the CNS. It is characterized by the destruction of myelin and myelin-producing oligodendrocytes. The disease etiology is associated with autoimmunity, pathogens such as viruses, and several genetic and environmental factors. Neuroinflammation in MS is mediated by the production of several inflammatory mediators, including cytokines, chemokines, and reactive oxygen and nitrogen species by activated glial cells, such as microglia and astrocytes, and CNS infiltrating myeloid and lymphoid cells [1, 2]. Reportedly. nitric oxide (NO) synthesized by inducible nitric oxide synthase (iNOS) significantly contributes to the oligodendrocyte degeneration, demyelination, and neuronal damage in MS [3].

iNOS metabolizes L-arginine to L-citrulline, generating NO as a byproduct in response to an inflammatory stimulus. Unlike its constitutively expressed isoforms eNOS (endothelial NOS) and nNOS (neuronal NOS), iNOS does not require calcium for its activity and, upon induction, produces NO in large amounts over extended periods [4]. It allows NO to interact with superoxide molecules to form peroxynitrites, a prominent marker of tissue injury and essential for pathogen elimination [5]. As iNOS is induced during inflammation, it is expressed by all leukocytes, including macrophages, dendritic cells, NK cells, primary tumor cells, and to a certain extent, activated T cells [6]. iNOS expression by microglia and macrophages is a prominent pro-inflammatory marker during neuroinflammation and is significantly implicated in the pathophysiology of MS [3].

MS patients revealed iNOS expression in the chronic active plaques in diverse cell types, including microglia/macrophages (MG/Mφ), ependymal cells, inflammatory cells, and to a certain extent in astrocytes [7]. Reactive astrocytes present in the acute MS lesions in the postmortem brains of MS patients were found reactive for iNOS mRNA and protein [8]. Results from experimental animal models of MS have shown mixed results. Several studies report improved disease symptoms upon inhibition of iNOS or iNOS deficiency [9-14], while in others, it rendered the system vulnerable to pathogen attack or exacerbated the disease [15-18]. Though studies implicating iNOS in demyelination are limited, most indicate a detrimental effect of iNOS, majorly because many reports are from the autoimmune model of MS where T cells and microglia and their interaction exert a pathogenic effect in the disease outcome.

In this study we delineate the role of iNOS in a murine β-coronavirus (MHV-A59) induced neuroinflammation model of MS wherein MG/Mφ is central to the pathogenesis and their interaction with T cells exhibit a protective role. Intracranial infection of demyelination susceptible C57BL/6 mice with hepato-neurotropic MHV-A59 or its spike gene recombinant strain RSA59 mimics the characteristic pathologies of MS and is one of the best-studied virus-induced models of MS [19]. A large repository of studies has revealed that RSA59 infection induces a biphasic disease in mice characterized by acute phase (day 5/6 post-infection [p.i.]) hepatitis, meningitis, encephalitis, and chronic phase (day 15-30 p.i.) demyelination and concomitant axonal loss [19-24]. RSA59 infection activates the CNS resident glial cells, including microglia and astrocytes, which mount the innate immune response [23, 24]. The immune cells of the brain further enhance inflammation by secreting an array of chemoattractants and pro-inflammatory cytokines, which facilitate peripheral leukocyte migration into the CNS [25]. Activated pro-inflammatory or M1-like microglia and other antigen presenting cells (APCs) present the viral antigens to the infiltrating T cells and stimulate the secretion of major antiviral cytokine IFNγ and other pro-inflammatory cytokines. Activated glial cells, along with the help of CD8+ and CD4+ T cells, clear the infectious viral particles and resolve the acute phase neuroinflammation as early as day 7 p.i. However, the viral mRNA persists in substantially low amounts through the chronic progressive demyelinating phase [26]. Anti-inflammatory cytokines secreted by CD4+ T cells induce a transition from pro to anti-inflammatory environment and reinstate the homeostasis in the CNS. The anti-inflammatory M2-like phagocytic microglia responsible for mopping the cell debris and initiating tissue regeneration harbor the demyelinating plaques in the spinal cord during the chronic phase. As a bystander effect, the chronically activated MG/Mφ are reported to strip the myelin from the axons resulting in demyelination, which starts as early as day 5-7 p.i. and reaches its peak at day 30 p.i. [26-29].

We previously showed an interplay between MG/Mφ and CD4+ T cells in the RSA59 induced neuroinflammation model wherein the absence of CD4+ T cells impaired viral clearance from the CNS and causes rare pathologies such as acute phase poliomyelitis, dorsal root ganglionic inflammation, and chronic phase axonal blebbing associated with severe demyelination. This study also reported that the exacerbated demyelination could result from persistent M2-like phagocytic MG/Mφ observed in the CD4-/- mice, indicating a protective role of CD4+ T cells [27]. Confirming evidence of the protective role of CD4+ T cells was further illustrated in a parallel study performed in Ifit2-/- mice. RSA59 infected Ifit2-/- mice presented with dampened activation of microglia/ macrophages resulting in low recruitment of CD4+ T cells in the CNS, which led to severe clinical disease and high mortality in the mice [28]. We also studied MG/Mφ interaction with CD4+ T cells on a molecular level with respect to CD40 ligand (CD40L), which is expressed by the activated CD4+ T cells and binds to CD40 receptor (CD40R) on MG/Mφ. Results showed that the absence of CD40L makes mice highly susceptible to RSA59 infection around day 10 p.i. This was attributed to the reduced activation and antigen presentation by microglia/ macrophage at the acute phase resulting in lower priming of CD4+ T cells in the cervical lymph nodes and reduced infiltration in the brain at day 10 p.i. Thus, leading to virus persistence and exacerbated chronic phase demyelination in the CD40L-/- mice mediated by phagocytic amoeboid microglia/ macrophages [29]. Studies have shown that CD40L binding to CD40R on microglia induces the production of iNOS [30]; we, therefore, wanted to examine the role of iNOS in RSA59 induced neuroinflammation comparing iNOS depleted mice in the background of demyelination susceptible C57BL/6 mice (iNOS-/-) and wild type C57BL/6 mice (WT) in this study to delve deeper into our understanding of CD4+ T cell and MG/Mφ crosstalk.

Here we show that RSA59 infection significantly upregulates iNOS expression during the acute phase (day 5 / 6 p.i.) when the neuroinflammation is heightened, and viral titer reaches its peak. In line with the earlier reports, iNOS expression upon RSA59 infection was significantly upregulated in immune cells of both myeloid and lymphoid origin, with macrophages, NK cells, and NKT cells being the major producers of iNOS. iNOS deficiency as assessed in 4 to 5 weeks old iNOS-/- mice compared to age-matched WT mice resulted in increased disease severity and mortality during day 9/10 p.i. which is the acute-adaptive transition phase. Infected iNOS-/- mice showed no difference in virus clearance compared to WT; however, increased infiltration of macrophages, neutrophils, and T regulatory cells were observed in the brains of iNOS-/- mice. The mRNA of TGFβ, a major anti-inflammatory cytokine and an inducer of T regulatory cells, was significantly upregulated in iNOS-/- mice. In addition, iNOS-/- mice, in comparison to WT mice, showed larger demyelinating plaques as early as day 9/10 p.i. populated with more amoeboid Iba1+ MG/Mφ. The marked presence of amoeboid MG/Mφ in the demyelinating plaques in iNOS-/- mice was further corroborated by the significant upregulation of the phagocytic marker CD206 and TREM2 and anti-inflammatory M2 marker Arginase 1 at mRNA level. Combining the results, our study highlighted that iNOS deficiency leads to the severity of demyelination at the acute-adaptive transition phase concurrent with a robust shift towards phagocytic MG/Mφ phenotype and alteration of the immune response towards an anti-inflammatory type in the RSA59 induced neuroinflammatory demyelination model.

## Results

### RSA59 infection induces iNOS expression in WT mice at the acute phase (day 5 p.i.)

Inflammatory stimuli are major factors that induce iNOS expression, which is otherwise strictly regulated within the cells [4]. Expression of iNOS transcripts upon RSA59 infection at 20000 PFUs was analyzed by quantitative real-time PCR of brain tissue from infected WT mice at days 6, 10, 15, and 30 p.i. The results showed significant upregulation of iNOS mRNA at the acute phase, i.e., day 6 p.i. (Figure 1 A). This coincides with the peak of the acute phase neuroinflammation in WT mice, which is characterized by elevated N gene mRNA levels, viral titer, and Iba1+ MG/Mφ activation as reported earlier [23, 31]. Flow cytometry further confirmed the upregulation in iNOS protein expression in brain from WT mice infected at 10000 PFUs at day 5 p.i. iNOS+ cells were gated from the live cell population, and their numbers and median fluorescent intensity (MFI) of iNOS expression were determined. The absolute numbers of iNOS expressing cells and the MFI for iNOS were significantly higher in the brain lysates of infected WT mice as compared to the mock infected controls (Figure 1 B, C, and D).

**Fig 1.**
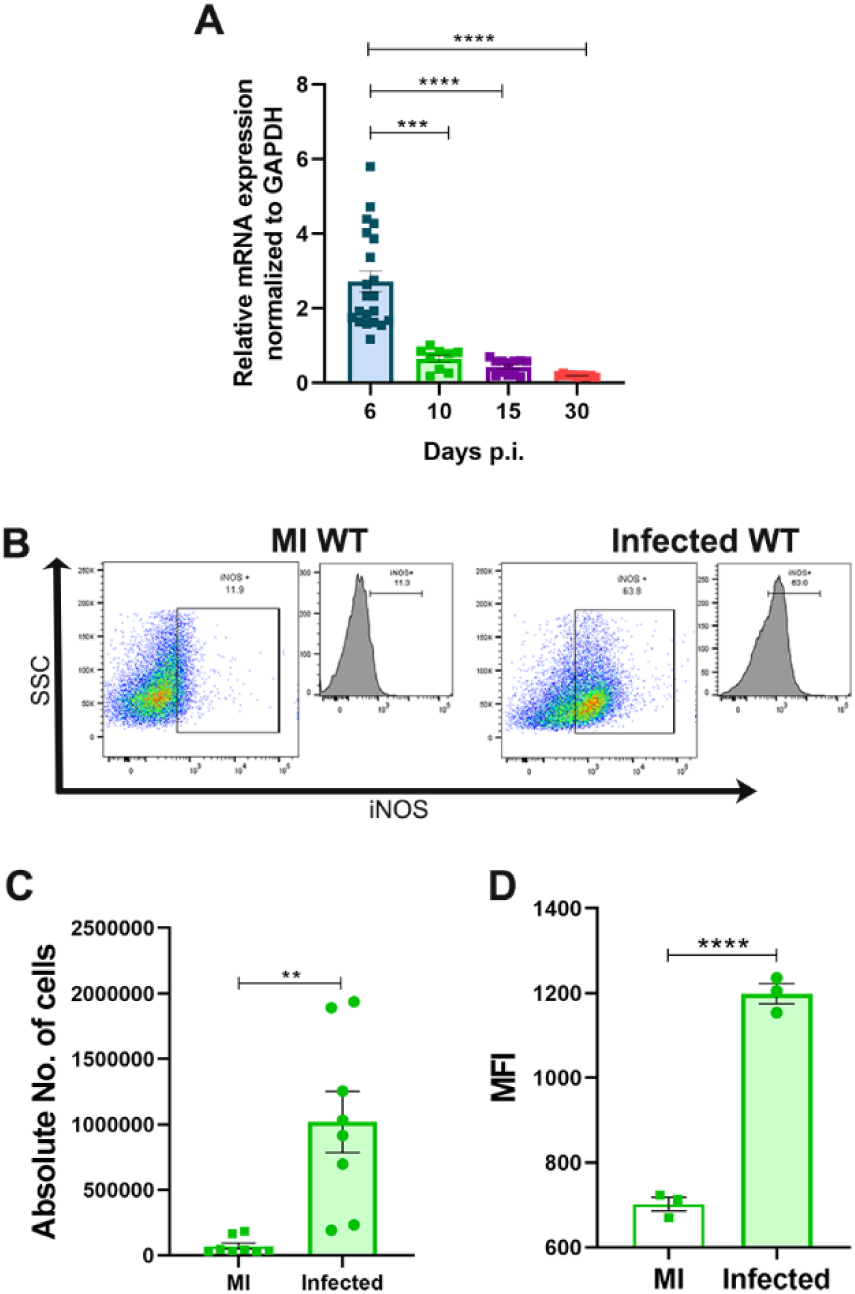
RSA59 infection upregulates the expression of iNOS in WT brains at the acute phase (day 5 p.i.). iNOS transcripts were determined in the brain of WT mice infected at 20000 PFUs at days 6, 10, 15 and 30 p.i. by qRT-PCR. Results were normalized to GAPDH compared and expressed relative to mock controls as Mean ±SEM (A). iNOS expression was determined by flow cytometry in the brain harvested from WT mice infected at 10000 PFUs at day 5 p.i. (B) shows representative flow cytometry dot plots and histograms showing percentages of iNOS+ cells, gated from live cell populations that were gated from singlets, and their absolute numbers and iNOS MFI are graphically represented in (C) and (D), respectively. *Asterisk represents statistical significance calculated using Kruskal-Wallis test for (A) and Welch’s t test for (C) and (D), p< 0.05 was considered as significant. *p< 0.05, **p< 0.01, ***p< 0.001, ****p< 0.0001. n= 3 to 7 mice per group.

### RSA59 infection in WT mice increases the expression of iNOS in macrophages, natural killer cells, and natural killer T cells

To further assess the source of iNOS at the acute phase, iNOS expression in different myeloid and lymphoid immune cell populations in the infected brains was determined by flow cytometry. Cells were gated from a parent gate of live iNOS+ cells gated from singlets, and the CNS resident and infiltrating myeloid cells were distinguished based upon CD45 low (CD45lo) and CD45 high (CD45hi) expression, respectively. The peripheral lymphoid populations were gated from iNOS+ CD3+ cells. Brain resident CD45lo CD11b+ microglia (Figure 2 A), CD45hi CD11c+ dendritic cells (Figure 2 B), CD45hi CD11b+ Ly6G-macrophages and CD45hi CD11b+ Ly6G+ neutrophils (Figure 2C), CD3+ CD4+ T cells (Figure 2 D), CD3+ CD8+ T cells (Figure 2 E), CD3+ NK 1.1+ natural killer T cells and CD3-NK1.1+ natural killer cells (Figure 2 F), showed increase in their absolute numbers of iNOS expressing cells on RSA59 infection at the acute phase. The iNOS MFI was upregulated in microglia, macrophages, CD8+ T cells, natural killer T cells, and natural killer cells (Figure 2A, C, E and F). However, there was no significant change in iNOS MFI in dendritic cells, neutrophils, and CD4+ T cells (Figure 2B, C and D). Further, it was observed that infiltrating macrophages, natural killer T cells, and natural killer cells showed comparatively high iNOS expression in the brains of the infected mice (Figure 2 G). In contrast, the MFI for iNOS was relatively high in CD8+ T cells, natural killer cells, and natural killer T cells compared to the myeloid cells (Figure 2 H/). Thus, at the acute phase of RSA59 infection, both myeloid and lymphoid immune cell subsets show an increase in the numbers of iNOS expressing cells with a maximum increase observed in macrophages, natural killer cell, and natural killer T cell population.

**Fig 2.**
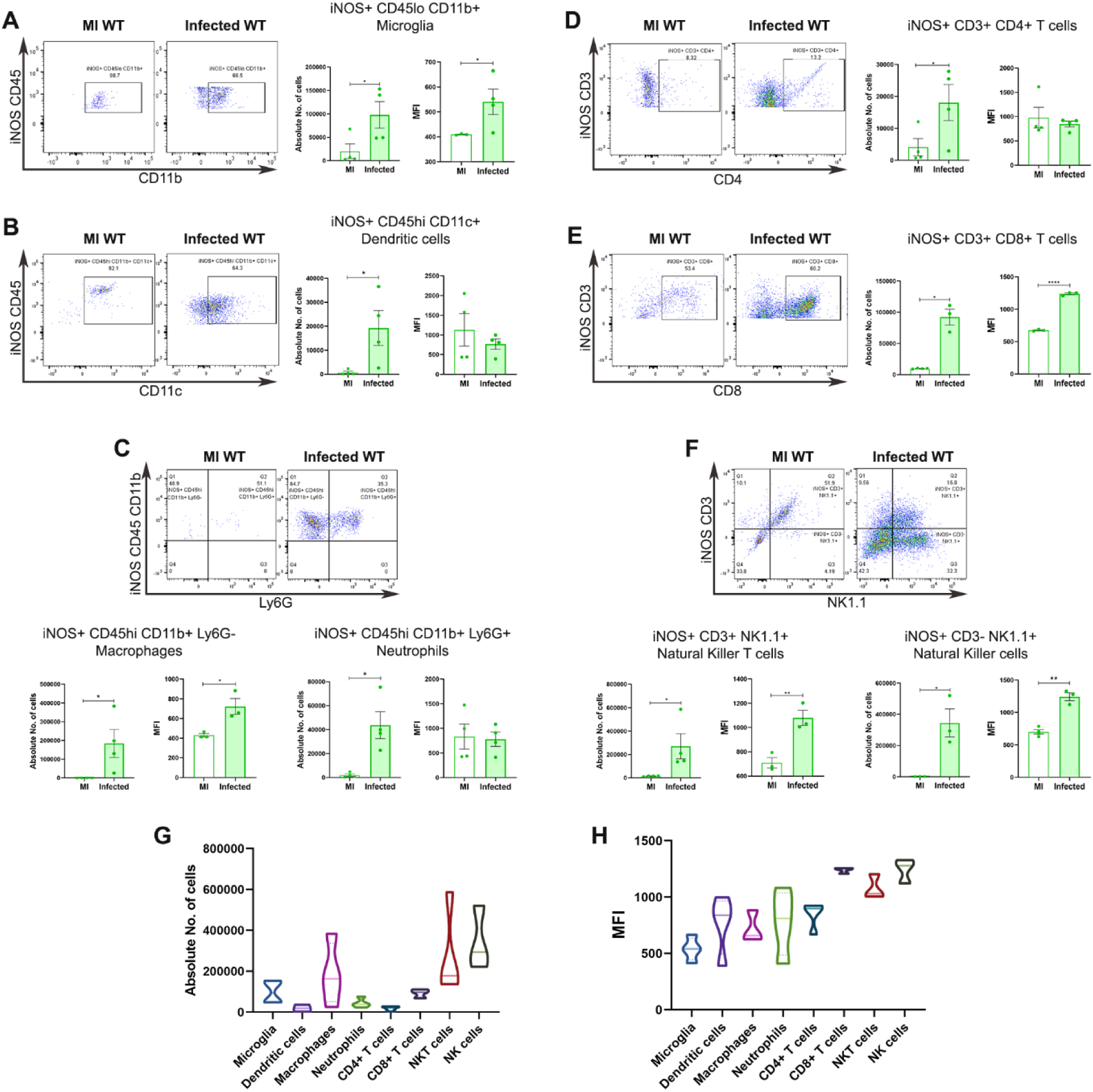
iNOS expression is significantly upregulated in myeloid and lymphoid immune cell subsets. Brains were harvested from mock and infected WT mice (10000 PFUs) on day 5 p.i. for flow cytometry analysis and stained for iNOS, CD45, CD11b, CD11c, Ly6G, CD3, CD4, CD8 and NK1.1. iNOS+ cells were gated from live cell populations gated from singlets. Subsequent immune cell subsets were gated from iNOS+ cells and their absolute numbers and iNOS MFI were compared between mock and infected WT groups. iNOS+ infiltrating myeloid cells were gated from iNOS+ CD45hi while iNOS+ resident microglia were gated from iNOS+ CD45lo. Lymphoid cell subsets expressing iNOS were gated from iNOS+ CD3+ for peripheral T cells and iNOS+ CD3-for NK cells. Representative dot plots and graphical representation of number of cells and iNOS MFI are given for iNOS+ CD45lo CD11b+ microglia (A), iNOS+ CD45hi CD11C+ dendritic cells (B), iNOS+ CD11b+ Ly6G-macrophages and iNOS+ CD11b+ Ly6G+ neutrophils (C), iNOS+ CD3+ CD4+ T cells (D), iNOS+ CD3+ CD8+ T cells (E), iNOS+ CD3+ NK1.1 (NKT) cells and iNOS+ CD3-NK1.1 (NK) cells (F). Volcano plot comparing iNOS+ cells in different cell subsets is represented in (G) and volcano plot comparing iNOS MFI in different cell subsets is represented in (H). Results were expressed as Mean ± SEM. *Asterisk represents statistical significance calculated using unpaired student’s t test with Welch’s correction, p< 0.05 was considered as significant. *p< 0.05, **p< 0.01, and ****p< 0.0001. n= 3 to 4 mice per group.

### iNOS deficiency results in heightened susceptibility to RSA59 infection

Mock infected WT and iNOS-/- mice did not show any differences in percent weight change up to day 10 p.i. indicating no inherent differences between the two strains (Figure S2 A). RSA59 infection at 20000-25000 PFUs in iNOS-/- mice resulted in more severe weight loss than WT infected mice (Figure 3 A). Clinical symptom assessment showed that the infected iNOS-/- mice developed the clinical disease as early as day 2 p.i. as opposed to beyond day 6 p.i. in infected WT mice. The clinical score reached up to 2 (characterized by ataxia, balance problem, and or partial paralysis) in iNOS-/- mice, while it did not increase beyond 1 (hunched back position with mild ataxia, possibly slower movement) in WT mice (Figure 3 B). Percent survival analysis showed that by day 10 p.i., only 72.22% infected iNOS-/- mice survived compared to the 100% of infected WT mice. Further, there was a drastic drop in survival around day 9/10 p.i., the acute-adaptive transition phase (Figure 3 C). Due to the low survival of the iNOS-/- mice infected with 20000 to 25000 PFUs of RSA59, some of the experiments were performed at 10000 PFUs.

**Fig 3.**
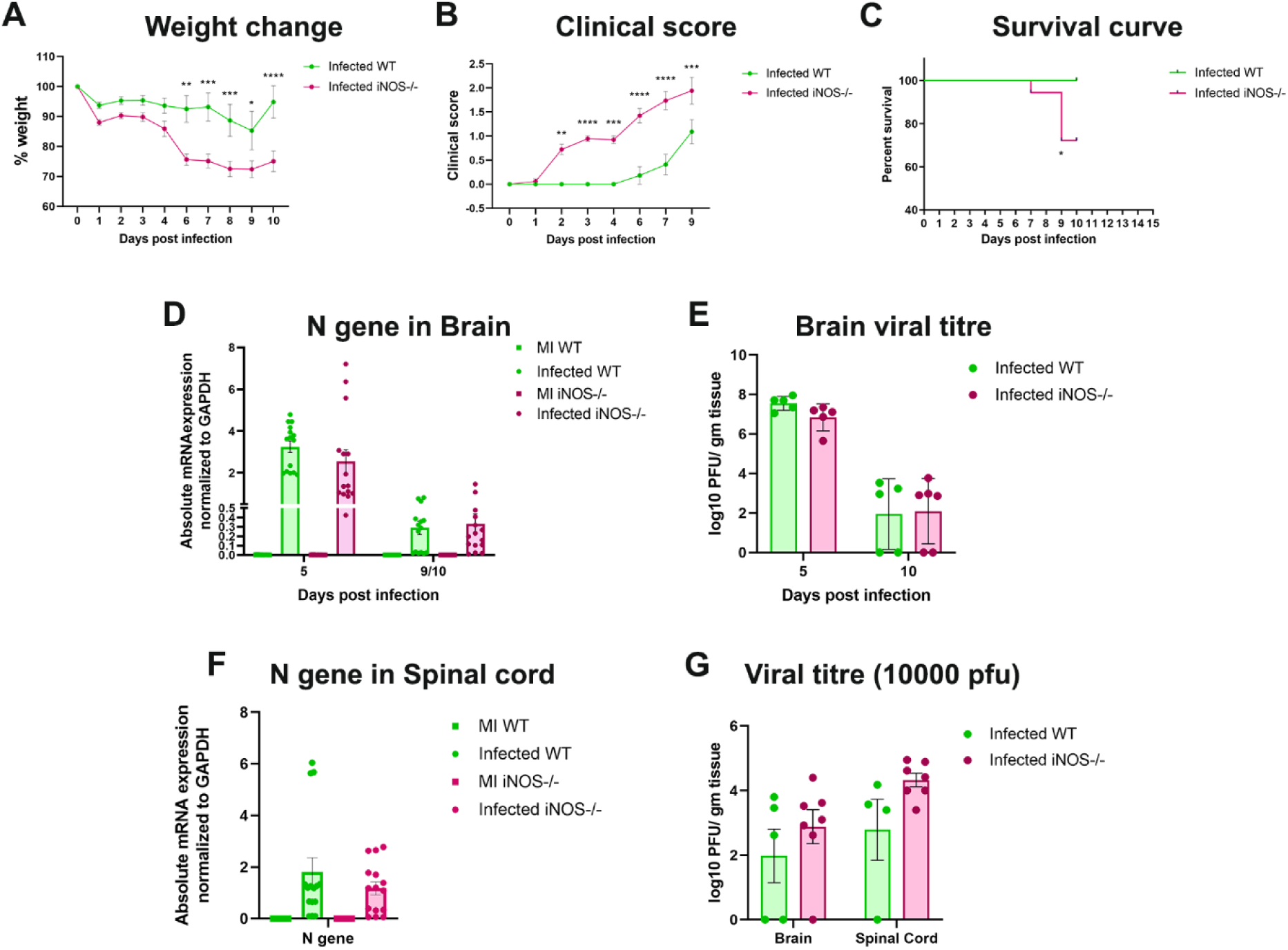
iNOS deficiency increases the disease severity in RSA59 infected mice but does not affect viral replication and persistence. Wildtype (WT) (n= 11 to 14) and iNOS-/-(n= 17 to 18) mice were infected with 20000 PFUs of RSA59 and monitored daily for weight change (A), development of clinical disease (B) and survival (C). Clinical scores were assigned by an arbitrary scale of 0–4 as described in Materials and Methods. WT is denoted by green color and iNOS-/- by pink. N gene (viral nucleocapsid gene) transcripts levels in brain tissue at days 5 and 9/10p.i. (D) and in spinal cord tissue lysate at day 9/10 p.i. (F) were determined by qRT-PCR and represented graphically as absolute mRNA expression normalized to GAPDH. (n= 3 to 5 mice per group per time point). Plaque assay was performed to determine viral titers in brains of mice infected at 20000 PFUs day 5 and 10 p.i. (E) and brains and spinal cord of mice infected at 10000 PFUs at day 9/10 p.i. (F). (n= 4 to 7 mice per group per time point). Statistical analysis was performed using Two-Way ANOVA with Tukey’s multiple comparison test for (A), (B), (D), (E), (F) and (G); and Log-rank (Mantel-Cox) test for survival proportions. Results were expressed as Mean ± SEM. *Asterisk represents statistical significance between infected WT and infected iNOS-/- groups. p< 0.05 was considered as significant. *p< 0.05, **p< 0.01, ***p< 0.001 and ****p< 0.0001.

### iNOS deficiency does not affect viral clearance by the system

Nitric oxide and iNOS are known to have antiviral activity [32]. We, therefore studied the viral load by quantifying the N gene mRNA and viral titers in the brains and spinal cords both at the acute phase, i.e., day 5 p.i., and at day 9/10 p.i. when the iNOS-/- mice showed a significant drop in their survival.

At both day 5 and day 9/10 p.i. WT and iNOS-/- mice infected with 20000 PFUs of RSA59 showed no differences in brain N gene transcript levels (Figure 3 D) and viral titers (Figure 3 E). No differences were observed in the N gene transcript levels of spinal cords in these mice at day 9/10 p.i. (Figure 3 F). Additionally, the viral titers in the brains and spinal cords of WT and iNOS-/- mice infected at 10000 PFUs at day 9/10 p.i. were comparable, implying no dose-dependent changes in the anti-viral response (Figure 3 G).

Together the data implied that iNOS deficiency did not affect the viral clearance in the CNS, and the disease severity observed in the infected iNOS-/- mice could be due to factors intrinsic to the immune responders.

### iNOS deficiency does not affect the brain pathology at both the acute and acute-adaptive transition phase

With differences in viral load in the CNS being ruled out as the cause of high mortality in infected iNOS-/- mice, the effect of iNOS on tissue pathology and MG/Mφ activation was examined next by H & E staining and immunohistochemical staining for the MG/Mφ activation marker, Iba1. Mock infected WT and iNOS-/- mice showed minimal to no inflammation in the brain, including Iba1+ MG/Mφ activation at both day 5 (Figure S1 B) and day 9/10 p.i. (Figure S2 C). Both WT and iNOS-/- groups infected with 20000 PFUs of RSA59 showed similar pathology on H & E staining with characteristic neuropathology of acute phase inflammation, i.e., meningitis (black arrow), vascular cuffing (green arrow), and microglial nodules (blue arrow) and equivalent staining for Iba1+ MG/Mφ at day 5 p.i. (Figure S3 A) Additionally, brains of infected WT and iNOS-/- mice at day 10 p.i. showed no differences in their pathology (Figure S3 C). Quantification of Iba1 expression in the infected brains at both day points further confirmed that iNOS deficiency did not affect MG/Mφ activation as evident by histopathological evaluation (Figure S3 B and D).

### iNOS deficiency results in increased infiltration of macrophages and neutrophils in the brain at the acute-adaptive transition phase

Further, the status of myeloid infiltration in the CNS at the acute-adaptive transition phase was investigated by flow cytometry in WT and iNOS-/- mice infected at 10000 PFUs. As expected, CD45hi peripheral immune cell numbers were significantly higher in both infected WT and iNOS-/- mice than their respective mock controls. In contrast, the CD45lo brain resident immune cell number did not vary across mock and infected groups (Figure 4 A). Interestingly, although no differences were found in Iba1+ MG/Mφ activation, the numbers of infiltrating CD45hi CD11b+ Ly6G-macrophages and CD45hi CD11b+ Ly6G+ neutrophils (Figure 4 C and E) were significantly higher in infected iNOS-/- mice compared to WT increased infiltration (Figure 4 B). CD45lo CD11b+ microglia numbers did not change in mock and infected WT and iNOS-/- mice (Figure 4 C).

**Fig 4.**
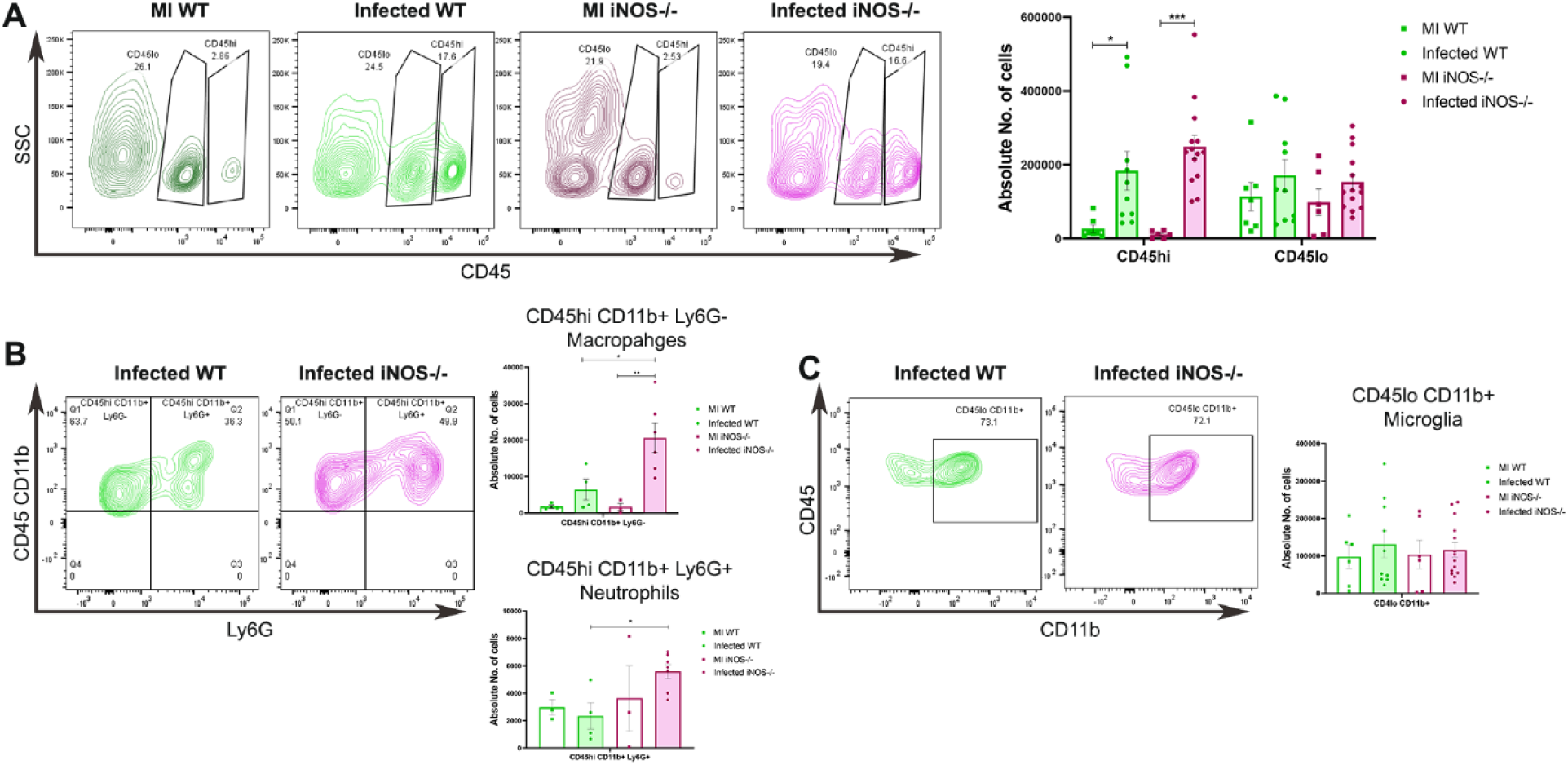
iNOS deficiency increases the infiltration of macrophages and neutrophils in RSA59 infected mice brains. Flow cytometry analysis was performed on brain from WT and iNOS-/- mock and RSA59 infected (10000 PFUs) brains at day 9/10 p.i. Infiltrating peripheral cells were distinguished by CD45hi gating from live cell populations gated from singlets. Similarly, CD45lo gate was used to extract brain resident cell populations. WT is denoted by green color and iNOS-/- by pink. Representative flow cytometry contour plots showing percentages of CD45hi and CD45lo populations and their graphical representation in mock and RSA59 infected WT and iNOS-/- groups are given in (A). Percentages of CD45hi CD11b+ Ly6G-macrophages and CD45hi CD11b+ Ly6G+ neutrophils are presented in representative contour plots and quantification of their numbers in graphical form in (B). (C) shows the representative contour plots of CD45lo CD11b+ microglia and its graphical representation of the absolute numbers. Statistical analysis was performed using Two-Way ANOVA with Sidak’s multiple comparison test between mock and infected WT and iNOS-/- groups. Results were expressed as Mean ± SEM. *Asterisk represents statistical significance. p<0.05 was considered as significant. *p< 0.05, **p< 0.01, and ***p< 0.001. n= 3 to 4 for MI and 5 to 7 for Infected.

### iNOS deficiency leads to a significant increase in the number of T regulatory cells and its inducer, TGFβ, in RSA59 infected brains at the acute-adaptive transition phase

Earlier, our studies have established the protective role of CD4+ T cells against RSA59 induced neuroinflammation and demyelination [27]. Whether iNOS can regulate peripheral lymphoid cell infiltration was studied next in WT and iNOS-/- mice infected at 10000 PFUs. No differences in the numbers of overall CD45hi CD3+ cells (Figure S4A), CD45hi CD3+ CD8+ T cells (Figure S4 B), CD45hi CD3+ NK1.1+ natural killer T cells and CD45hi CD3-NK1.1+ natural killer cells (Figure S4 C) were observed. Also, the total CD45hi CD4+ CD4+ T cell population showed no differences between infected WT and iNOS-/- brains (Figure 5 A). But CD45hi CD3+ CD4+ CD25+ Foxp3+ T regulatory subpopulation numbers were significantly higher in the infected iNOS-/- mice (Figure 5 B). Additionally, mRNA levels of TGFβ, one of the major inducers of T regulatory cells and a potent anti-inflammatory marker, were also significantly upregulated (Figure 5 C).

**Fig 5.**
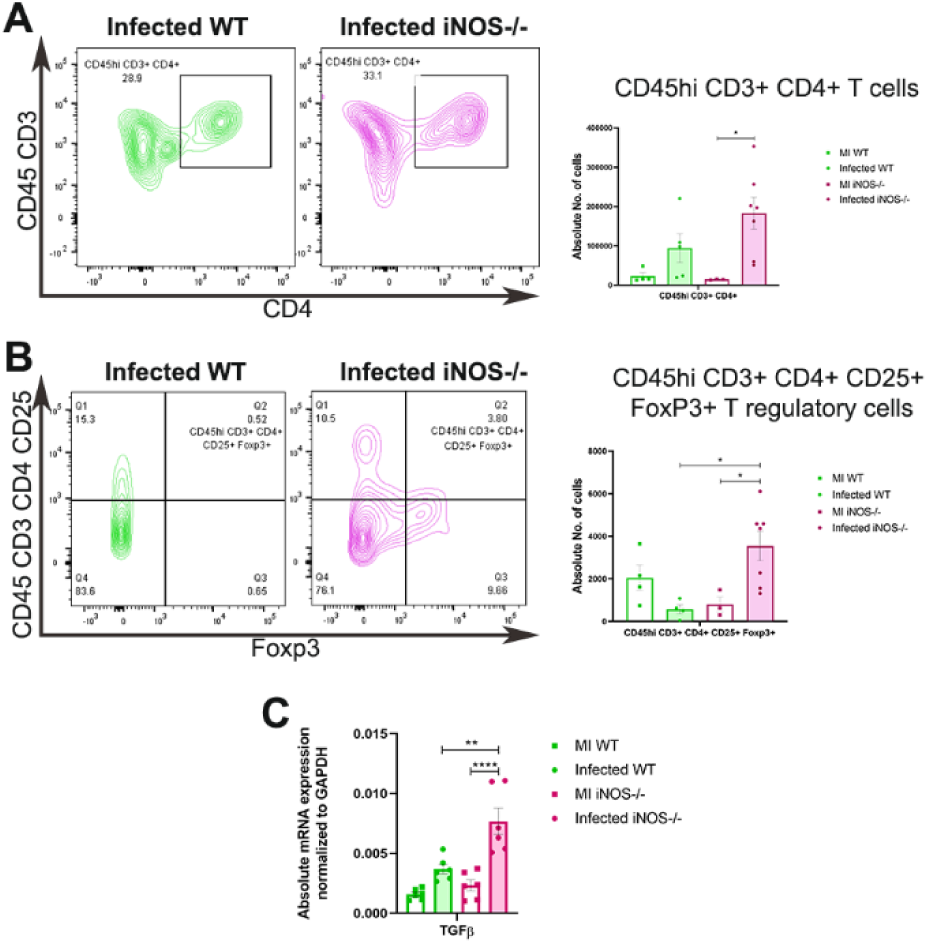
iNOS deficiency leads to increased infiltration of T regulatory cells and upregulation of TGFβ mRNA expression in the brain of RSA59 infected mice at the acute-adaptive transition phase (day 9/10 p.i.). Brains were isolated from mock and infected (10000 PFUs) mice brains at day 9/10 p.i. and flow cytometry analysis was performed to check lymphoid cell subsets. Peripheral T cells were gated from CD45hi CD3+ cell populations extracted from live cells gated from singlets. WT is denoted by green color and iNOS-/- by pink. (A) shows representative contour plots of CD45hi CD3+ CD4+ T cell percentages and graphical representation of their absolute numbers. Regulatory CD4+ T cells were gated by CD45hi CD3+ CD4+ CD25+ FoxP3+ markers and their representative contour plots are shown in with quantification of absolute numbers of Tregs are represented in (B). The mRNA levels of TGFβ were determined by qRT-PCR from brain of mock and infected WT and iNOS-/- brains and are represented as absolute mRNA expression normalized to GAPDH in (C). Statistical analysis was performed using Two-Way ANOVA with Sidak’s multiple comparison test between mock and infected WT and iNOS-/- groups. Results were expressed as Mean ± SEM. *Asterisk represents statistical significance. p< 0.05 was considered as significant.*p< 0.05, **p< 0.01, and ****p< 0.001. n= 3 to 4 for MI and 5 to 7 for Infected.

### iNOS deficiency enhances demyelination and leads to an increased accumulation of amoeboid MG/Mφ in the spinal cords of RSA59 infected mice at the acute-adaptive transition phase

Previous studies demonstrated that RSA59 induced demyelination starts as early as day 5 p.i. and reaches its peak at day 30 p.i. [19]. The status of spinal cord inflammation on both day 5 and day 9/10 p.i. and demyelination at day 9/10 p.i. was evaluated in WT and iNOS-/- mice infected at 20000 PFUs.

H & E staining and immunohistochemical staining of Iba1 in mock infected spinal cords at day 5 p.i. (Figure S1 C) and H & E staining, Luxol fast blue staining, and immunohistochemical staining of Iba1 at day 9/10 p.i. (Figure S2 D) revealed no disease phenotypes. Infected WT and iNOS-/- mice at day 5 p.i. exhibited similar pathology (Figure S5 A) and levels of Iba1+ MG/Mφ activation (Figure S5 B).

However, striking differences were observed in the spinal cord pathology of infected WT and iNOS-/- mice at day 9/10 p.i. H & E staining of the spinal cord sections showed higher myelitis in the spinal cords of infected iNOS-/- mice than WT mice (Figure 6 A, black arrows). In addition to this, Luxol fast blue staining (LFB) revealed increased demyelination in the iNOS-/- mice (Figure 6 B, blue arrows). The demyelinating plaques had a characteristic accumulation of Iba1+ MG/Mφ. Quantification of the percent demyelination and Iba1 expression in the spinal cords of infected mice revealed significantly high demyelination in the absence of iNOS (Figure 6 D) but a similar expression of Iba1 by the activated MG/Mφ (Figure 6 E). Both infected WT and iNOS-/- showed the presence of activated ramified MG/Mφ in the grey matter (Figure 6 C, black-lined rectangles), but the MG/Mφ in the white matter exhibited notably different morphologies (Figure 6 C, brown lined rectangles). The MG/Mφ in the white matter of WT mice were similar to those in their grey matter with few interspaced amoeboid cells. On the contrary, the white matter MG/Mφ in the demyelinating plaques of the iNOS-/- mice were characteristically amoeboid in morphology, denoting a phagocytic phenotype.

**Fig 6.**
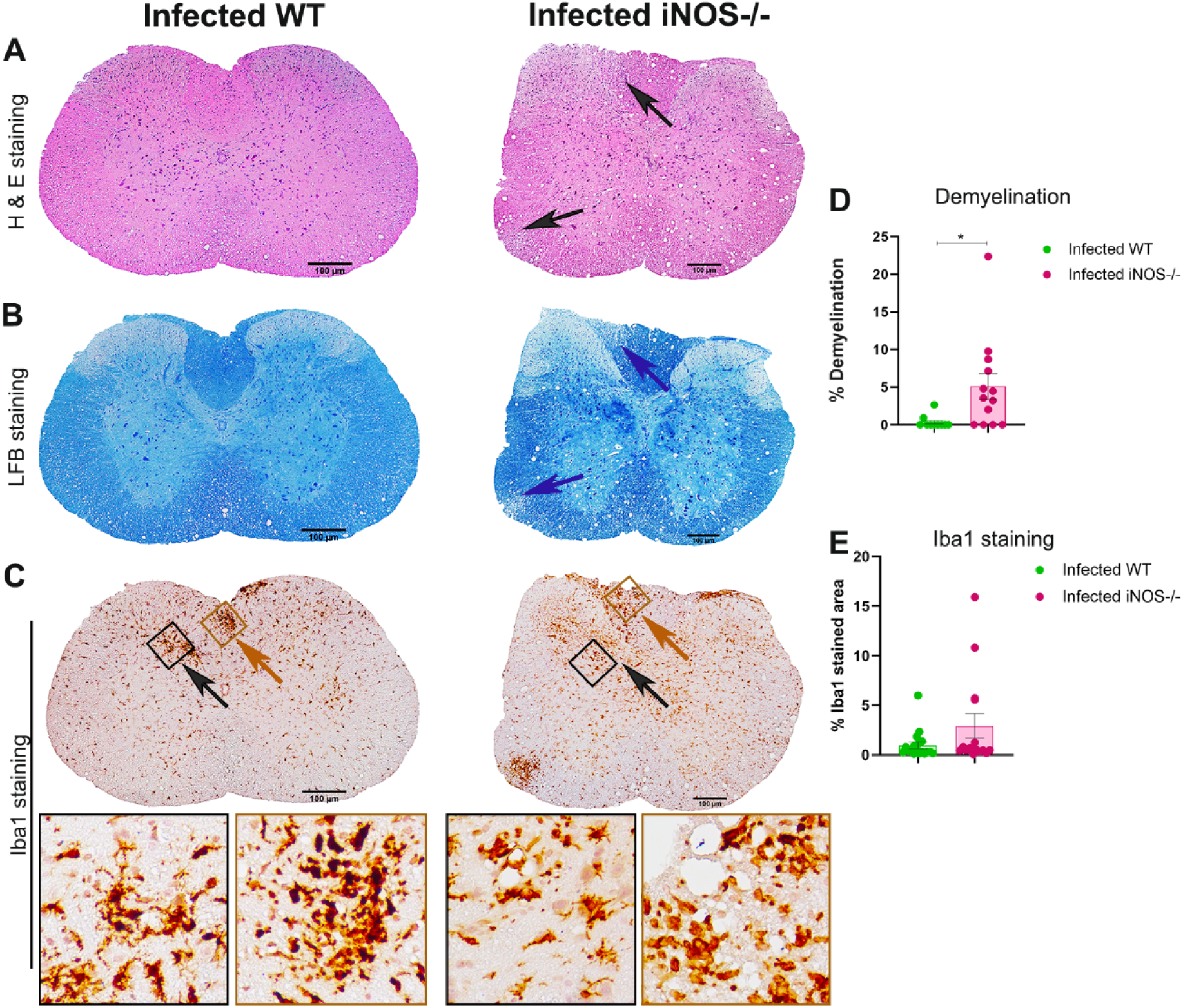
iNOS deficiency led to increased demyelination with markedly high presence of amoeboid microglia/ macrophages in the spinal cords of RSA59 infected mice at the acute-adaptive transition phase (day 9/10 p.i.).

5µm thick cross-sections of RSA59 infected (20000 PFUs) WT and iNOS-/- mice spinal cords were stained for the presence of inflammatory lesions by H & E (A), demyelination by LFB (B) and activated MG/Mφ by Iba1 (C). Black boxed areas highlighted by black arrow represent higher magnification of grey matter area, and brown boxed areas highlighted by brown arrow represent higher magnification of white matter below the corresponding Iba1 stained spinal cord cross-sections. Scale bar 100µm. (D) and (E) are the quantification of percent demyelination and percent Iba1 stained area, respectively. WT is denoted by green color and iNOS-/- by pink. Results were expressed as Mean ± SEM. *Asterisk represents statistical significance calculated between infected WT and infected iNOS-/- mice using unpaired student’s t test with Welch’s correction. p< 0.05 was considered as significant. *p< 0.05. n= 4 to 6 mice per group.

### iNOS deficiency leads to upregulation of phagocytic and anti-inflammatory markers characteristic of M2-like phenotype at the acute-adaptive transition phase

The amoeboid morphology of MG/Mφ is associated with an anti-inflammatory M2 phagocytic phenotype [33]. To verify the MG/Mφ phenotype status, the transcript levels of different phagocytic markers were assessed at day 9/10 p.i. Results showed that microglial phagocytic marker TREM2 and M2 marker Arg1, the antagonist of iNOS, were significantly upregulated in both the brains (Figure 7 C and D) and spinal cords of iNOS-/- mice (Figure 7 G and H). Another phagocytic marker, CD206, showed significant upregulation in the spinal cords of iNOS-/- infected mice (Figure 7 F) but not in the brain (Figure 7 A). In contrast, P2Y6 did not change in either of the infected groups (Figure 7 B and F).

**Fig 7.**
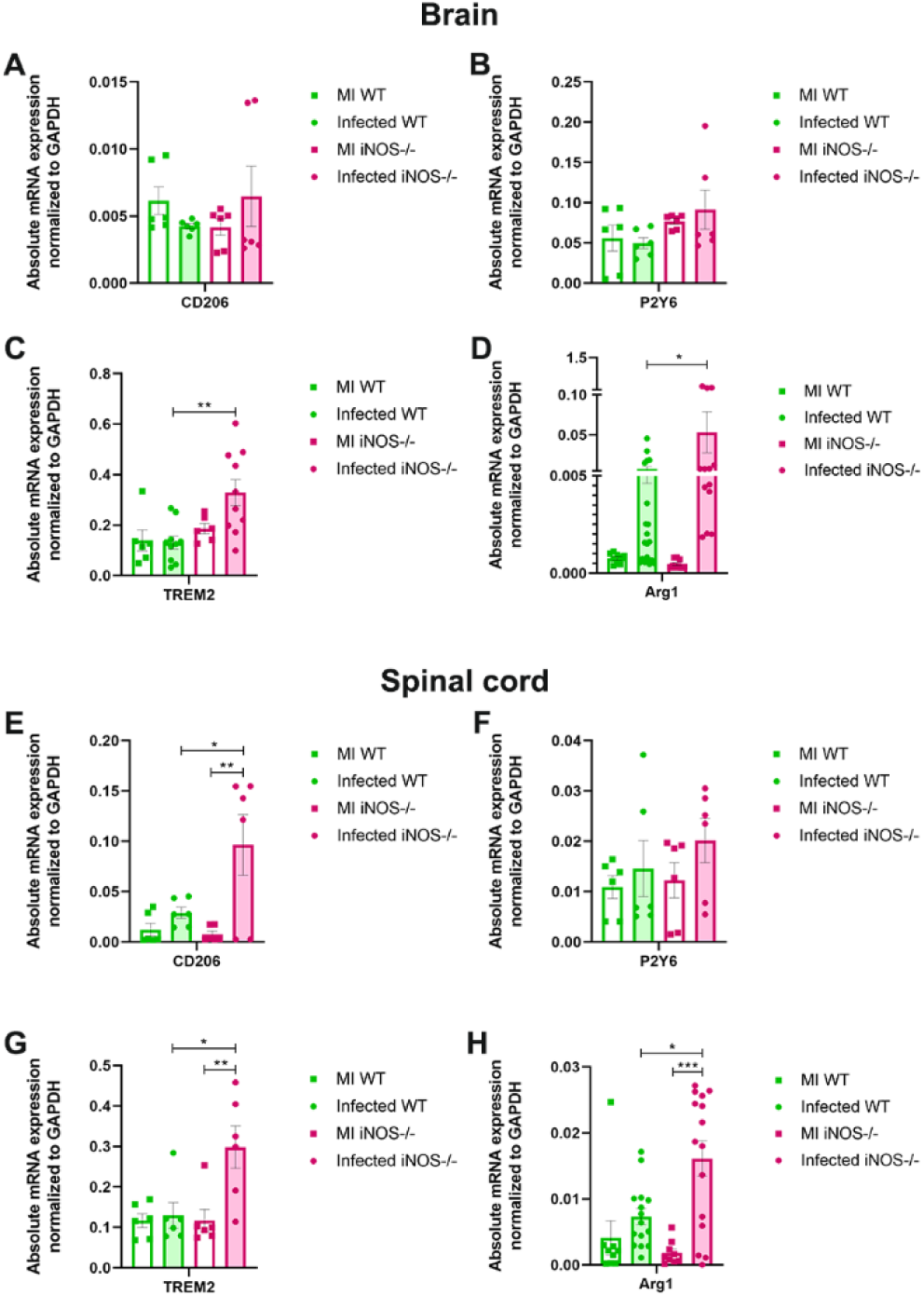
iNOS deficiency resulted in upregulation of anti- inflammatory and phagocytic markers of microglia/ macrophages in RSA59 infected mice brain and spinal cords at day the acute-adaptive transition phase (day 9/10 p.i.). mRNA was isolated from brain and spinal cord tissue of mock and infected (20000 PFUs) RSA59 WT and iNOS-/- mice. Transcript levels of CD206 (A and E), P2Y6 (B and F), TREM2 (C and G) and Arg1 (D and H) were analyzed by qRT-PCR. WT is denoted by green color and iNOS-/- by pink. Results were expressed as Mean ± SEM. *Asterisk represents statistical significance using Two-ANOVA with Tukey’s multiple comparisons test. p< 0.05 was considered as significant. *p< 0.05, *p< 0.05, **p< 0.01, and ***p< 0.001. n= 3.

Thus, although pathologically no differences were observed in Iba1+ activation of brain MG/Mφ, the mRNA levels of few M2-like phenotype markers were upregulated in both the brain and spinal cords of iNOS-/- mice. In conclusion, in the absence of iNOS, the MG/Mφ showed an aggravated anti-inflammatory phagocytic phenotype, which may contribute to increased demyelination and, therefore, high disease severity in iNOS-/- mice.

## Discussion

In this study, we showed that intracranial infection of RSA59 in wildtype mice led to a significant increase in the expression of iNOS in myeloid and lymphoid cell subsets in the brain during the acute phase of the disease with maximum iNOS expression found in macrophages, natural killer cells, and natural killer T cells. Studies carried out in iNOS-/- mice showed that despite no significant differences in the viral clearance by the CNS, the absence of iNOS leads to heightened disease severity and mortality. Further, iNOS-/- mice showed a significant increase in the infiltration of macrophages and neutrophils in the brain at day 9/10 p.i. i.e., the acute-adaptive transition phase. This phase was also marked by increased numbers of Tregs and expression of its inducer TGFβ in iNOS-/- mice as compared to the WT mice. Severe demyelination was observed in the spinal cords of these mice concurrent with the presence of amoeboid microglia/ macrophages. The absence of iNOS led to significant upregulation of TREM2 in the brain and CD206 and TREM2 in spinal cords. The signature M2 marker Arg1 also showed a significant increase in both brain and spinal cords of iNOS-/- mice at the acute-adaptive transition phase. Combined, these results revealed a role of iNOS in regulating a timely transition towards an anti-inflammatory CNS phenotype and thus preventing the development of severe demyelination at an early phase.

The role of iNOS in neurodegeneration has been studied in murine experimental models of MS utilizing demyelination inducing chemicals, peptides, or virus strains, the outcomes of which often depend on differential disease pathology and the role of infiltrating T cells. Studies in MHV strain V5A13.1 (MHV-V5A13.1) induced demyelinating disease reported that NO generated by iNOS did not affect the viral clearance and the demyelination status but resulted in increased neuronal apoptosis [13]. Another study incorporating a highly neurovirulent strain MHV-JHM showed that iNOS deficiency or inhibition did not affect MHV demyelination [34]. Studies in the neurotoxicant cuprizone-induced demyelination model, on the other hand, showed that the absence of iNOS exacerbated demyelination at early time points [17]. We have previously reported the central role of microglia/macrophage in demyelination and the protective role of CD4+ T cells and its ligand CD40L through its interaction with microglia/macrophage in MHV-RSA59 induced neuroinflammatory disease [27, 29]. iNOS has been reported to be induced in BV-2 microglial cells on ligation of CD40 [30]. MHV-RSA59 infection in C57BL/6 WT mice led to upregulation of iNOS at the acute phase, i.e., day 5 p.i. in different immune cell subsets, including macrophages, microglia, dendritic cells, CD4+ T cells, CD8+ T cells, natural killer T cells, and natural killer cells. iNOS deficiency in these mice led to increased mortality and early demyelination. However, it did not affect viral clearance, which is in line with the previous reports. We also observed that iNOS deficiency led to increased infiltration of macrophages and neutrophils in the RSA59 infected brains at day 9/10 p.i. i.e., acute-adaptive transition phase, although the activation states of MG/Mφ did not differ compared to the WT mice. Slightly similar alteration in myeloid cell infiltration was reported in a relatively recent study in the experimental autoimmune encephalomyelitis (EAE) model of MS where it was showed that inhibition of iNOS with L-NAME altered the brain pathology at the antigen-priming phase while the spinal cord pathology showed differences in the effector phase. Additionally, there was high recruitment of pathogenic CD11b+F4/80−Gr-1+ cells and low infiltration of regulatory CD11b+F4/80− cells in the brain during the priming phase and in the spinal cord during the effector phase [35]. CXCL12 is a known chemokine that limits CNS cell infiltration. A study in rats immunized for EAE reported that increased iNOS expression reduced CXCL12 gene expression in spinal cord homogenates. The expression was brought back to normal with inhibition of nitric generation *in vivo* [36]. The effect of iNOS in chemokine production by glial and CNS infiltrating cells warrants further investigation in RSA59 induced disease model.

CD4+ T cells, in previously published reports by our group, have shown to play a protective role in RSA59 induced demyelination [27]. In this study, we did not find any change in the numbers of total infiltrated CD4+ T cells in the brain. However, the numbers of CD4+ CD25+ FoxP3+ T regulatory cells were almost double in infected iNOS deficient mice brains. Further, the transcript levels of TGFβ, a potent pleiotropic anti-inflammatory marker and an inducer of Tregs [37, 38], were significantly upregulated as well. Tregs are known to exert homeostatic effect by suppressing T effector cells and have also been extensively implicated in promoting oligodendrocyte health, regeneration, and remyelination [39, 40]. A groundbreaking study by the Fitzgerald group in the mouse model of MS clearly showed the involvement of Tregs in remyelination [41]. However, in our study, we found that the iNOS deficient RSA59 infected mice exhibited more Tregs and early severe demyelination at the acute-adaptive transition phase. Since CD4+ T cells play a protective role in RSA59 model of MS, significantly more demyelination in infected iNOS deficient mice could be due to suppression of T cell function by the Tregs. A study in Theiler’s murine encephalomyelitis virus (TMEV) infection model of MS showed that introducing *ex vivo* generated induced Tregs on day 0 magnified the acute phase disease. This was partly due to the suppression of the antiviral response by the effector T cells. The viral titers were thus high and CNS recruitment of peripheral cells low [42]. Additionally, a study in murine syngeneic tumor model reported inhibition of Treg induction by iNOS expressed by CD4+ T cells. iNOS knockout or inhibition, in turn, resulted in Treg induction [43]. Studies in mice deficient in CD4+ T cells, Ifit2 or CD40L have shown that the CD4+ T cells are protective and crucial for MG/Mφ homeostasis during RSA59 infection [27-29]. Though the virus clearance remains unaffected, whether the increase in the numbers of Tregs affects the functions of effector T cells in infected iNOS deficient mice remains to be checked.

Histopathology study of spinal cord tissues at day 9/10 p.i. revealed the marked presence of amoeboid MG/Mφ in the demyelinating lesions of iNOS deficient mice. This was accompanied by significant upregulation of M2 MG/Mφ phenotype markers such as CD206 and Arg1 in the brain and CD206, TREM2, and Arg1 in spinal cords, confirming the anti-inflammatory phagocytic phenotype of MG/Mφ in iNOS deficient mice [44, 45]. Earlier studies have reported that overexpression of TREM2 in microglia led to enhanced phagocytosis and reduced expression of pro-inflammatory cytokines while its knockdown upregulated iNOS [45]. Arginase 1 is the most prominent maker of alternatively activated M2 macrophages and competes with iNOS for the same substrate, L-arginine [46]. MG/Mφ in the tissue do not exhibit any one phenotype state but rather tune their polarization in response to the inflammatory milieu. Even so, the expression of iNOS and Arg1 is considered a signature marker of pro-inflammatory (classically activated M1-like) and anti-inflammatory (classically activated M2-like) phenotypes. A lucid study in the EAE induced model of MS targeted the expression of iNOS and Arg1 by mononuclear phagocytes entering the brain and studied the journey of these cells in the inflamed CNS. The cells specify and adapt to their phenotype locally guided by the CNS-derived signals and can change their individual phenotypes over time accordingly [47].

iNOS deficiency leads to enhanced anti-inflammatory or M2-like MG/Mφ phenotype, which may be responsible for exacerbated demyelination in RSA59 infected mice. It is important to note that CD4+ CD25+ FoxP3+ Tregs can assert their effects on monocyte/ macrophages and drive their polarization towards the M2 state [48, 49]. Also, although TGFβ is considered essentially neuroprotective [50-52], its high local expression in the CNS was shown to increase the infiltration of cells in the CNS, leading to CNS impairment and faster disease onset in an EAE model of MS [53]. Further, a study reported that TGFβ inhibition downregulates Arg1 expression in primary microglia [54]. Arg1 can suppress effector T cells by limiting L-arginine [55, 56]. Thus, in a system such as RSA59 infected CNS, where CD4+ T cell function is required to maintain tissue homeostasis by suppressing MG/Mφ activation, enhancement of anti-inflammatory phagocytic functions of MG/Mφ and CD4+ T cell inhibiting environment might result in sustained MG/Mφ activation states leading to tissue damage.

Over the past years, studies in experimental models of MS have shown that not anyone but all the phases of the immune response, innate, adaptive, and the bridging of the aforementioned, are crucial and thereby determinants of the outcome of the disease [57, 58]. The majority of the immune responders identified to play a role in these various phases during the course of MS do not operate in a mutually exclusive manner but rather synergistically, and the resultant of these interactions may vary based on the system used. The timescale of these responses is critical and can alter the usually beneficial effects of the immune responders. We showed that the absence of one of the significant MG/Mφ pro-inflammatory markers skews the trajectory of the immune response towards the anti-inflammatory type and heightens the disease severity leading to early demyelination. iNOS can potentially affect the infiltration of cells in the CNS and the polarization states of MG/Mφ and is thus crucial in regulating the timely transition of the system from pro to anti-inflammatory status. The nexus between iNOS, Arg1, and T regulatory cells may open a new avenue in understanding the underlying mechanics of pro to anti-inflammatory switch, which will aid in designing therapeutics targeted at ameliorating tissue damage associated with neuroinflammatory demyelination. These mechanisms can be further extended to explore the pathology of neurological manifestations in SARS-CoV-2 owing to the similarities in beta coronavirus pathogenesis.

## Materials and methods

### Ethics statement

iNOS-/- mice (Jackson Laboratory, B6.129P2-Nos2tm1Lau/J, Stock no. Stock No. 002609) [59] were obtained from the Animal Facility at the National Centre of Cell Science Pune, India. All experimental procedures and animal care and use were strictly regulated and reviewed per animal ethics approved by the Institutional Animal Care and Use Committee at the Indian Institute of Science Education and Research Kolkata (AUP no. IISERK/IAEC/AP/2019/45). Experiments were performed following the guidelines of the Committee for the Purpose of Control and Supervision of Experiments on Animals (CPCSEA), India.

### Virus, inoculation of mice, and experimental design

Mice were infected with RSA59, an isogenic recombinant strain of the demyelinating mouse hepatitis virus; MHV-A59 described previously [60]. Five-week-old MHV- free wild type (WT) B6 mice (Jackson Laboratory) and iNOS-/- (B6.129P2-Nos2tm1Lau/J) mice (Jackson Laboratory; Stock No. 002609) were used in the study. The mice were inoculated intracranially with RSA59 at 20000 (50% of 50% lethal dose, LD50) and 10000 PFUs. Similarly, mock infected controls were inoculated with phosphate-buffered saline (PBS) plus 0.75% bovine serum albumin (BSA). Mice were monitored daily post-infection for disease signs and symptoms, and changes in their weight were noted. Clinical disease severity was graded using the following scale: 0: no disease symptoms; 0.5: ruffled fur; 1: hunched back with mild ataxia; 1.5: hunched back with mild ataxia and hind limb weakness; 2: ataxia, balance problems and or partial paralysis; 2.5: one leg completely paralyzed, motility issues but still able to move around with difficulty; 3: severe hunching/wasting/paralysis of both hind limbs and severely compromised mobility; 3.5: severe distress, complete paralysis, and moribund; 4: dead. For RNA studies, mice were sacrificed on day 6 (acute phase), day 10 (acute-adaptive transition phase), day 15 (chronic adaptive phase), and day 30 p.i. (chronic phase). For histopathological, immunohistochemical analyses, viral titer estimation, and flow cytometry experiments, mice were sacrificed at day 5 and 9/10 p.i.

### Estimation of Viral Replication

WT and iNOS-/- mice infected with RSA59 at 10000 PFUs and 20000 PFUs were sacrificed at day 5 and 9/10 p.i. Brains were harvested and placed into 1 ml of isotonic saline containing 0.167% gelatin (gel saline). Brain tissues were weighed and kept frozen at -80°C until titered. Tissues were subsequently homogenized and using the supernatant, viral titers were quantified by standard plaque assay protocol on tight monolayers of L2 cells as described previously [60] using the formula: plaque-forming units (PFUs) = (no. of plaques X dilution factor/ml/gram of tissue) and expressed as log10 PFUs/gram of tissue.

### Flow Cytometry Analysis

Mice were transcardially perfused with 20ml PBS, and half brains were homogenized in 2 ml of RPMI containing 25 mM HEPES (pH 7.2), using Tenbroeck tissue homogenizers. Tissue homogenate was then centrifuged at 450g for 10 min. Obtained cell pellets were resuspended in RPMI containing 25 mM HEPES, adjusted to 30% Percoll (Sigma), and underlaid with 1 ml of 70% Percoll. A second centrifugation cycle was applied at 800 g for 30 minutes at 4°C, following which the cells were recovered from the 30%-70% interface, washed with RPMI, and suspended in FACS buffer (0.5% bovine serum albumin in Dulbecco’s PBS). Specific cell types were identified by staining with fluorochromes like fluorescein isothiocyanate (FITC), phycoerythrin [(PE), (PECy7)], peridinin chlorophyll protein [(PerCP), (PerCpCy5.5)], allophycocyanin [(APC), (APCCy7)] and violet excitable dyes [(V450), (V500)] conjugated MAb for 45 minutes on ice in FACS buffer. Expression of surface markers was characterized with MAb (all from BD Biosciences except where otherwise indicated) specific for CD45 (clone Ly-5), CD11b (clone M1/70), CD11c (clone HL3), CD3 (clone 145-2C11), CD4 (clone GK1.5), CD8 (clone 53-6.7), NK1.1 (clone PK136), CD25 (clone PC61), and Foxp3 (clone G155-178). Intra-nuclear staining was performed for Foxp3 after fixation and permeabilization (BD 562574) per the manufacturer’s guidelines.

Samples were acquired on a BD FACSVerse flow cytometer (BD Biosciences) and analyzed on FlowJo 10 software (Treestar, Inc., Ashland, OR) [28]. First, doublet exclusion using FSC-A and FSC-W was performed, and then cells were gated based on forward scatter (FSC), and side scatter (SSC) to focus on live cells. Further, the cells were analyzed to differentiate myeloid and lymphoid populations. Myeloid cells were gated from a primary gating on CD45, and an independent CD3 gating or CD3 gating in addition to CD45 was applied for the lymphoid populations. Single colors and FMOs were used in all the experiments. Beads were gated based on their FSC/SSC pattern.

### Histopathology and immunohistochemical analysis

Mice were sacrificed at day 5 and day 9/10 p.i. Following transcardial perfusion with PBS and post-fixation with 4% paraformaldehyde for 26-48 hours, liver, brain, and spinal cord tissues were harvested and embedded in paraffin. Five-micrometer-thick sections of the embedded tissues were prepared and stained with hematoxylin and eosin for histopathologic analysis. Luxol fast blue (LFB) staining was performed to evaluate demyelination in the spinal cord tissues, as described previously with minor modifications [27]. MG/Mφ activation in the brain and spinal cord sections was analyzed using 1:500 dilution of polyclonal antibody directed against MG/Mφ-specific calcium-binding protein; Ionized calcium-binding adaptor molecule 1 (Iba1) (Wako; Catalog No. 019-19741). Bound primary antibodies were detected by an avidin-biotin immunoperoxidase technique (Vector Laboratories) using 3,3-diaminobenzidine (DAB) as the substrate [61]. Control slides from mock infected mice were stained in parallel. All the slides were coded and read in a blinded manner.

### Quantification of histopathological sections

Image analysis was performed using Fiji’s basic densitometric thresholding application (Image J, NIH Image, and Scion Image) as described previously [62]. Briefly, image analysis for Iba1 stained sections was performed by capturing the images at the ×4 for brain and ×10 for the spinal cord so that the entire section (i.e., the scan area) could be visualized within a single frame. The RGB image was deconvoluted into three different colors to separate and subtract the DAB-specific staining from the background hematoxylin staining. The perimeter of each brain and spinal cord tissue was digitally outlined, and the area was calculated in square micrometers. A threshold value was fixed for each image to ensure that all antibody-marked cells were taken into consideration. The amounts of Iba1 staining were termed the percent area of staining.

To determine the area of demyelination, LFB stained spinal cord cross-sections from each mouse were chosen and analyzed using Fiji (Image J, NIH Image, and Scion Image) [31, 62]. The total perimeter of the white matter regions in each cross-section was marked, and the area was calculated by adding together the dorsal, ventral, and anterior white matter areas in each section. Also, the total area of the demyelinated regions was outlined and collated for each section separately. The percentage of spinal cord demyelination per section per mouse was calculated.

### RNA isolation, reverse transcription, and quantitative PCR

RNA was extracted from brain tissues of RSA59 infected WT and iNOS-/- mice and mock infected mice using the TRIzol isolation protocol following transcardial perfusion with diethylpyrocarbonate (DEPC)-treated PBS. The total RNA concentration was measured using a NanoDrop ND-2000 spectrophotometer. One microgram of RNA was used to prepare cDNA using a high-capacity cDNA reverse transcription kit (Applied Biosystems). Quantitative real-time PCR analysis was performed using a DyNAmo ColorFlash SYBR Green qPCR kit (Thermo Scientific) in a Step One Plus real-time PCR system (Thermo Fisher Scientific) under the following conditions: initial denaturation at 95°C for 7 min, 40 cycles of 95°C for 10 s and 60°C for 30 s and melting curve analysis at 60°C for 30 s. Reactions were performed in duplicates or triplicates. Sequences for the primers used are given in Table 1. Absolute quantitation was achieved using the threshold (ΔCT) method. mRNA expression levels of target genes in mock and RSA59 infected WT and iNOS-/- mice were normalized with GAPDH and expressed as absolute mRNA expression to depict the changes between the infected WT and iNOS-/- groups as well as between the mock infected if any.

**Table 1.**
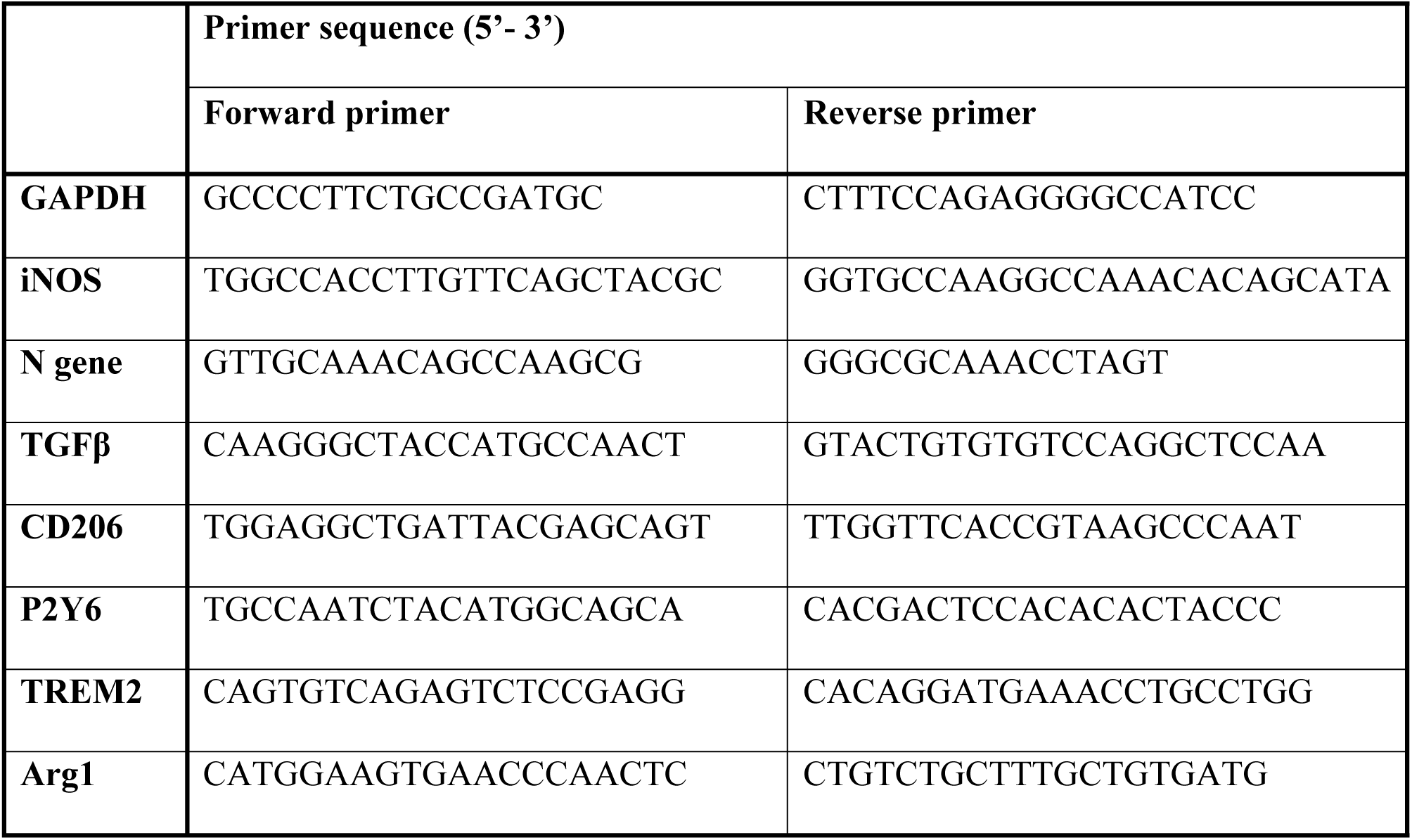
List of primers used for quantitative real time PCR.

### Statistical analyses

Values were represented as mean values ± standard errors of the mean (SEM). Values were subjected to unpaired student’s t-tests with Welch’s correction or Kruskal-Wallis One-Way ANOVA or Two-Way ANOVA with multiple comparison tests (Tukey’s test and the Holm-Sidak test) for calculating the significance of differences between the means. Log-rank (Mantel-Cox) test was used for calculating the statistical significance in mortality between groups. All statistical analyses were done using GraphPad Prism 8 (La Jolla, CA). A P-value of <0.05 was considered statistically significant.

## Availability of data and materials

This study includes no data deposited in external repositories. All relevant data are within the manuscript and its supporting information files.

## Consent for publication

All authors have read the final manuscript and approved it for publication.

## Author contributions

MK and JDS designed and planned all the experiments. SK and FS performed flow cytometry experiments along with MK and helped analyze the results. MK and JDS analyzed the data and wrote the manuscript. FS blindly quantified histopathological data, assigned clinical scores to the experimental mice, and helped MK write the manuscript. BS helped in critically reviewing the manuscript. MK, FS, and JDS participated in data interpretation. JDS supervised and reviewed the overall study.

## Acknowledgments

We thank the Animal facility at the National Centre of Cell Science, Pune, India, for providing the iNOS-/- mice (B6.129P2-Nos2tm1Lau/J, Jackson Laboratory; Stock No. 002609) used in the study. We thank the IISER-Kolkata animal facility for providing the necessary support. We thank DBT for supporting and funding the Flow cytometry facility.

